# Novel Natural Product Discovery from Marine Sponges and their Obligate Symbiotic Organisms

**DOI:** 10.1101/005454

**Authors:** Regina R. Monaco, Rena F. Quinlan

## Abstract

Discovery of novel natural products is an accepted method for the elucidation of pharmacologically active molecules and drug leads. Best known sources for such discovery have been terrestrial plants and microbes, accounting for about 85% of the approved natural products in pharmaceutical use (1), and about 60% of approved pharmaceuticals and new drug applications annually (2). Discovery in the marine environment has lagged due to the difficulty of exploration in this ecological niche. Exploration began in earnest in the 1950’s, after technological advances such as scuba diving allowed collection of marine organisms, primarily at a depth to about 15m.

Natural products from filter feeding marine invertebrates and in particular, sponges, have proven to be a rich source of structurally unique pharmacologically active compounds, with over 16,000 molecules isolated thus far (3, 1) and a continuing pace of discovery at hundreds of novel bioactive molecules per year. All classes of pharmaceuticals have been represented in this discovery process, including antiprotazoals, pesticides, TGF-beta inhibitors, cationic channel blockers, anticancer, cytotoxic, antiviral, anti-inflammatory and antibacterial compounds. Important biosynthetic pathways found in sponges which give rise to these compounds include the terpenoid (4), fatty acid, polyketoid, quinone reductase, alkaloid, isoprenoid (5), and non-ribosomal protein synthase pathways.

## 1. Introduction

### 1.1

Humans have long sought medicines, antidotes, palliatives, recreational substances and other bioactive materials that enhance and/or positively affect health from their environment. From early medicinal treatments such as willow bark tea for pain or extracts of foxglove plants for cardiomyopathy, to the far more efficacious current-day pharmaceuticals which include cholesterol lowering statins, anti-cancer drugs such as doxorubicin, bleomycin, taxol and rapamycin, narcotic analgesics such as the opioids and non-narcotic analgesics such as ziconitide, or antibiotics such as penicillin, the cephalosporins, tetracycline and its derivatives – more than 60% of modern pharmaceuticals are directly derived from natural product research (1, 2). Research and successful development of pharmaceutical agents continues to remind us that the natural world remains an incompletely mined source of novel pharmaceutical agents (6). As for the de-novo synthetic design of pharmaceutical agents, it may be commented that human ingenuity alone could not have led to the design of most of the known, naturally derived pharmaceutical agents or leads. This observation is based on comparison of the chemical structures of completely synthetically designed pharmaceutical agents as compared to the chemical structures of natural products.

The investigation of micro-environments found within rainforests, deserts, and even urban sewers, has yielded novel molecular products from organisms that populate those environments. Promising novel lead compounds possess pharmaceutical properties unknown before the natural world yielded them up to further investigation. While most drug discovery has resulted from the investigation of environmental niches within forests, soils and lakes, there has been less investigation of the marine environment to about 15 meters in depth, and virtually no investigation at greater depths. It is important to recognize that the exploration and harvesting that has already occurred in the terrestrial niches has been somewhat exhaustive, and many drug leads are actually being “rediscovered”, a process which wastes precious research resources. The marine environment is a nearly unexplored resource, and is reasonably expected to yield many novel compounds.

The marine environment has been less effectively mined to date than terrestrial environments due to its inherent inaccessibility, and the difficulty of preserving and transporting samples from their origin to the laboratory. The advent of technological advances, including and beginning with scuba diving in the 1950s and continuing to date, as well as advances in preservation of biological samples, has accelerated discovery. It has already been shown that this environment is a rich arena for pharmaceutical discovery. The marine environment represents a unique evolutionary niche yielding unique bioactive molecules. This is due in part to the nature of the environment itself, as well as many marine organisms’ obligatorily evolved arsenal of competitive molecular survival tools within that environment. Most of these protective molecules are secondary metabolites - biologically active molecules not directly involved in the normal functions of the organism, functions which include growth, reproduction or development. Secondary metabolites are dynamically synthesized by the organism in direct response to environmental stressors and thus act as protective and/or adaptive agents. Such responses include up-regulation of toxic compounds as protection, or in response to damage or predator attack; adaption to heat, mechanical deformation, salinity, pH, or cold; aggression (stinging) or other environmental stressors such as UV irradiation, overgrowth, competition for mates, tissue damage, local lack of essential nutrients. Secondary metabolite concentration within an organisms’ cells is thus variable and environmentally responsive.

Marine organisms have evolved to produce distinct and dynamic molecular responses due to the stressors in their environment, such as nearly ubiquitous bacterial and viral competition and attack, and continuously variable environmental parameters such as salinity or pH. Additionally, marine organisms tend to share a higher degree of networked interconnectedness than land organisms, and do not suffer from degenerative diseases similar to mammalian life. As example, aging mechanisms differ or are absent (i.e., no telomere shortening is observed), and cancer and cardiovascular disease (or more accurately, its equivalent) are unobserved. Marine organisms generally exist symbiotically, they communicate with external, diffusible pheromones, some have the ability to create camouflage in unique modalities (both internally and externally), and they may metabolically utilize minerals such as silica and/or rare, toxic metals such as vanadium or gold via biochemical pathways which are not found in land-creatures.

Thus, the marine ecosphere may be considered a unique evolutionary niche which has given rise to a dynamic, diverse mélange of novel molecular structures generated by the force of evolution, a self-adaptive and organizing force in a dynamic universe. Exploration of such evolutionary niches using methods of genomic data mining has and will continue to yield novel potential pharmaceutical agents.

### 1.2 Sponge Biology

Sponges (phylum: Porifera) are nearly ubiquitous invertebrate sedentary filter-feeders, found abundantly throughout the marine environment and in freshwater, with about 10,000 known species thriving from the polar oceans to tropical oceans and seas. Their structure is that of an asymmetric collection of multiple types of specialized eukaryotic cells, with no organs, circulatory, nervous or digestive systems. A gelatinous mesogleal matrix is held together by mineralized and/or fibrous skeletal elements (observed as “spicules”). The organization of the cells within the mesoglea is highly canalized with a single layer of flagellated cells, known as choanocytes, lining the pores. Seawater is driven through the pores, nutrients extracted, and cellular waste is transported outward through the interior canals by the choanocytes, exiting through the single osculum pore of the animal. This action captures a diverse array of bacteria in addition to serving the purpose of filtering for food particles.

Although a few species each of carnivorous and solitary sponges exist, most sponges symbiotically inhabit coral reefs, where they participate in nutrient cycles with organisms in the coral reef. Endosymbiotic fungi, bacteria and viruses extensively and obligately co-habitate with marine sponges. These organisms participate in the nutrient cycle of the sponge as well as in the production of the protective, chemically unusual secondary metabolites. It is these bioactive molecules that are the primary subject of natural product discovery in sponges. It should be noted that most of these microorganisms are unculturable [79], with current discovery taking place via manual washing of the sponge and careful handling of sponge tissue and wash liquid to capture the organisms that inhabit sponge cells directly.

This internal microenvironment may be thought of as loosely similar to that of “microbial mats” found in rainforests. Although lacking the structure and integrity of terrestrial microbial mats, endosymbioic microflora of sponges most likely inhabit internal domains of the sponge depending on their function – for example, photosynthetic organisms will live near the outer surface of the animal, where they can collect the most light.

### 1.3. Discovery

Traditionally, natural product discovery involves standard protocols of screening, extraction and identification. To enhance this methodology and add to the efficiency of the discovery process, genomic searching is used. This became a powerful discovery tool beginning in the 1990’s as both computational power and advances in machine learning led to new or increasingly improved methods to search genomic data. Techniques such as sequence alignment combined with available structural information of proteins involved in pathways known to lead to secondary metabolite synthesis, such as the polyketoid or terpenoid pathways, lead to elucidation of the biochemical steps along these metabolic pathways of interest and/or to candidate molecules expected to possess pharmacological activity.

Genomic searching techniques use known sequences of genes that have been determined to be involved in secondary metabolite metabolic pathways, the most-studied gene clusters for this being the hybrid polyketide and non-ribosomal peptide synthases (PK-NRPS) [43], the isoprenoid synthases, and the terpenoid synthases. Each of these synthase genes represents a multi-functional gene cluster, which yields secondary metabolites that may be tested for activity. Note that, as exemplified in the polyketide synthase (PKS) gene cluster, this same gene cluster will yield substantially different secondary metabolites, characteristic of the particular organism, due to the ability of PKS to exist with its components (which are each single-function metabolic enzymes) in differing, evolutionarily shuffled orders. The gene product is metabolized along the PKS gene cluster in a linear fashion, thus the order of the subunit genes results in an organism-specific secondary metabolite. This is why PKS, and similar multi-component gene complexes, are such rich targets for genomic discovery.

Thus, genomic searches are carried out by searching a genomic database for sequence motifs that indicate their origin from known biosynthetic pathways such as PKS. An example of this is the isoprenoid pathway, which is the pathway preceding carotenoid synthesis. In this case one would start with a novel natural product that has shown some structural or sequence similarity to known genes along the isoprenoid pathway in genomic database. It may be noted that lycopene synthase is the committed entry point for metabolism of the carotenoids. Also note that terpenes are molecular subunits involved in the synthesis of a wide variety of other biological products, not just the carotenoids.

Modern bioinformatic computational methods are far more powerful discovery tools than the traditional pharmaceutical discovery methodology of “grind and find.” The existing set of bioinformatic software for genomic, metabolomic, proteomic, and related “omics” database mining is being constantly developed, improved and optimized, and a large variety of software to carry out sophisticated, compute-intensive searches such as alignment of multiple genomes or comparison of multiple sequences or sequence targets exists and is often freely available.

The use of computational genomics allows the researcher to design computational searches targeted to discover particular classes or types of novel natural products. This may be done in new as well as previously studied genomes. In the computational searching of genomes that have already been studied using traditional wet lab methodology, additional discovery of bioactive novel products that had been overlooked often occurs. Also, such computational database searches are more thorough than ever before, due to rapidly improving and increasingly efficient software tools utilizing “big data” algorithmic approaches, and machine learning techniques, which can direct and optimize which search paths to explore, in ways that human researchers might not choose unaided.

After discovery of promising novel natural products, candidate molecules are further studied computationally, using computational chemistry techniques including molecular dynamics (MD), normal modes analysis (NMA), and energy minimization. Derivatives of these promising lead compounds, which may be chemically modified at reactive moieties via, as example, glycosylation, methylation, epimerization, dimerization, reduction, or oxidation, are studied. A semi-synthetic library of enhanced candidate molecules is compiled and analyzed, and drug screening is carried out to discover if enhanced activity of the lead compound via such functional substitution occurs.

Mass spectroscopy (MS) in conjunction with computational genomics is used for the characterization of newly discovered natural products. Peptide products may have their sequence identified using MS and then be associated with their gene of origin from their unmodified sequence [15]. Natural product peptidogenomics (NPP) sequence tagging is done to determine the amino acid sequence, which is translated into the genetic sequence and used to search for the originating gene or gene cluster in the genome or metabolome being studied. QqQ mass spectrometry methods are becoming increasingly quantitative in achieving absolute quantification of low-concentration metabolites that are difficult to detect using NMR.

Computational genomic software can also search, not only for molecular motifs, but also for sequences and sequence similarities. For example, NP.searcher [16] searches genomic databases and returns predicted gene clusters, while MORPH [17] searches genomic data using sets of known genes from a chosen metabolic pathway, and returns all associated genes, thus building an information base of the organism’s metabolic pathways and overall metabolic network organization. NP.searcher may also return a linear peptide sequence if a candidate gene is identified. Note that the software will generate many such molecular sequences as the genome is searched which must be further evaluated. Sequence alignment of multiple genomes allows comparative searching and is expected to discover similar candidates over different genomes, and will construct best-guess molecule products for each genome.

### 1.4 Examples of bioactive secondary metabolites and pharmaceutical agents which have been discovered from marine sponges

#### i. Bioactive Secondary Metabolites

##### Terpenoids

Terpenoids (or isoprenoids), which include both primary and secondary metabolites, are a large and diverse class of compounds derived from C_5_ isoprene units [18, 19]. Of the plethora of natural products found in marine organisms, the terpenoids are especially prominent and widespread. Terpenes, and other products of secondary metabolism in marine organisms such as sponges, play a vital ecological role. Sponges are sessile and soft-bodied, thus they have developed potent bioactive secondary metabolite defensive compounds [20]. Natural products found in or associated with sponges have pharmaceutical relevance due to their antitumor, antiviral, antimicrobial, and antiprotozoal properties [21, 22].

**Figure 1.**
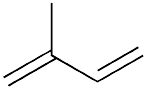
Structure of Isoprene (C_5_H_8_) [19x].

##### Sesterterpenoids (C_25_)

Sesterterpenoids (C_25_) represent a major class of marine terpenoids identified in sponges, and the bioactivities and pharmaceutical activities of these compounds have been well-studied and characterized [23] Manoalide is a potent antibiotic and the parent compound of a series of pharmaceutically active sesterterpene metabolites isolated from the marine sponge *Luffariella variabilis* [24]. These compounds were shown to have antimicrobial activity against Gram-positive bacteria including *Staphylococcus aureus* and *Bacillus subtilis* [25] A class of bicyclic sesterterpenoids with anti-tumor activity is the thorectandrols, isolated from the marine sponge *Thorectandra*. All thorectandrols, including the parent compound of the group palauolide [26] and palauolol [27] have been assayed for antiproliferative activity against human tumor cell lines and found to be active against all but one of the cell lines assayed [28].

Sponges of the genus *Hippospongia* are a source of various sesterterpenoids, which possess a broad spectrum of bioactivities, including isocitrate lyase (ICL) inhibitory [29], cytotoxic [30-35], antispasmodic [36], and antibacterial [37] activities. Recently Chang et al., (2012) isolated two novel sesterterpenoids from *Hippospongia* sp. - the pentacyclic sesterterpene, hippospongide A, and a scalarene sesterterpenoid, hippospongide B [38]. The sesterterpenes heteronemin [39], heteronemin acetate [40] and hyrtiosin E [41] exhibited significant cytotoxicity against human colon adenocarcinoma, hormone-dependent breast cancer, and human chronic myelogenous leukemia cell lines [38].

##### Triterpenoids (C_30_)

Another class of terpenoids of frequent occurrence in marine sponges are the triterpenoids (C_30_). Two highly bioactive triterpenoidal metabolite families are the isomalabaricane triterpenes, and the steroidal saponins, both of which have been isolated from marine sponges [42]. The isolation of isomalabaricane triterpenes was first reported from the Fijian sponge *Jaspis stellifera* [42] and the Somalian sponge *Stelleta* sp. [43].

Isomalabaricane triterpenoids. Central to this class is the yellow triterpenoidal stelletin A [44], isolated from J. stellifera [42] and found to be cytotoxic against a murine leukemia cell line (P388) exhibiting an IC50 of 2.1 nM [85].

**Figure 2.**
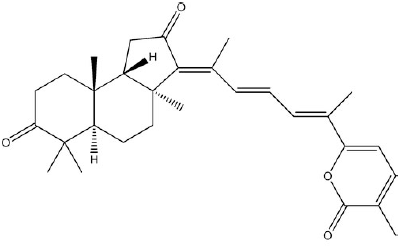
Structure of Stelletin A [84].

A second major group of isomalabaricane triterpenes are the stelliferins, which includes 13 known compounds. Among these, stelliferins A-F, derived from J. stellifera, show significant antineoplastic activity against both murine lymphoma (L1210) (IC50 of 1.1-5.0µM) and human epidermoid carcinoma (KB) cells (IC50 of 2.8-13.0µM) [86]. The potent antiproliferative activities exhibited by stelletins and stelliferins has led to the discovery of chemical synthetic methods for both stelletins and stelliferins [46].

The isomalabaricane triterpenoids globostellatic acids, first isolated from the marine sponge *Stelletta globostellata*, have cytotoxicity profiles similar to that of the stelletins and stelliferins. These compounds also demonstrate potent cytotoxicity against murine leukemia P388 cells [47]. In addition, globostelletins A-I, along with other isomalabaricane-derived natural products including jaspolides F and rhabdastrellic acid-A from *Rhabdastrella globostellata*, were also shown to induce inhibitory activities in human gastric gland carcinoma, human intestinal adenocarcinoma and human hepatocellular carcinoma cells. Furthermore, rhabdastrellic acid-A was demonstrated to induce apoptosis of human promyelocytic leukemia (HL-60) cells [48].

##### Steroidal saponins

Steroidal and triterpene glycosides are some of the more pharmaceutically relevant metabolites isolated from marine sponges. The acidic steroidal metabolite penasterol (a close relative of lanosterol) was isolated from the Okinawan sponge *Penares* sp. and has been shown to possess potent antileukemic activity [49].

Other steroidal saponins with important bioactivities include the erylosides that have been reported from different species of the genus Erylus. The first eryloside congener isolated from the Red Sea sponge Erylus lendenfeldi was Eryloside A, which has demonstrated significant antitumor activity against murine leukemia P388 cells, as well as antifungal activity against Candida albicans. Erylosides E and F, isolated from, respectively, the Atlantic sponge E. goffrilleri [50] and E. formosus, revealed immunosuppressive and potent thrombin receptor antagonistic activities [51].

##### Other pharmaceutically relevant classes of terpenoids in sponges

Biosynthetically, meroterpenoids are a group of mixed polyketide and terpenes, typically isolated from fungi and marine organisms. Several novel meroterpenoids, including a new structural group of meroterpenoid metabolites, the insuetolides A-C, as well as several drimane sesquiterpenes, were recently isolated from the marine-derived fungus *Aspergillus insuetus* (OY-207), which was isolated from the Mediterranean sponge *Psammoncinia* sp. collected off the coast of Israel. A number of these new secondary metabolites were shown to have important pharmacological acitivites. Insuetolide A exhibited anti-fungal activity, and both insuetolide C, as well as the new drimane sesquiterpene, (E)-6-(4′-hydroxy-2′-butenoyl)-strobilactone A, demonstrated mild cytoxicity against MOLT-4 human leukemia cells [52].

Thirteen terpenoids isolated from *Spongia* sp. and *Ircinia* sp., collected from the Turkish coastline of the Aegean Sea [53] were recently reported to exhibit potent antiprotozoal activity. The linear meroterpene 4-hydroxy-3-tetraprenylphenylacetic acid demonstrated activity against the parasitic protozoan *Typanosoma brucei rhodesiense*, a causative agent of African typanosomiasis (sleeping sickness), without any cytotoxicity. In addition, Orhan et al. [52] reported that the diterpenoid dorisenone D exhibited the best antiplasmodial efficacy against the malarial parasite *Plasmodium falciparum*, with some activity against *T. brucei rhodensiense*.

A large number of novel secondary metabolites exhibiting a diverse array of both biological activities and chemical structures have been derived from the *Clathria* genus of marine sponges. The isolation of three new bicyclic C_21_ terpenoids, clathric acid, two derivatives of N-acyl taurine, and clathrimides A and B from the sponge *Clathria compressa* were recently reported. Antimicrobial assays show that clathric acid possesses mild antibacterial activity [54].

##### Carotenoids (C_40_)

Carotenoids are C_40_ terpenoids synthesized by tail-to-tail linkage of two molecules of C_20_ geranylgeranyl diphosphate. The polyene skeleton of carotenoids is the most structurally distinguishing feature, and consists of a long system of alternating conjugated double and single bonds in which the π-electrons are delocalized along the entire length of the polyene chain; this feature is responsible for the characteristic molecular shape, light-absorbing properties, and chemical reactivity of carotenoids [55].

**Figure 3.**
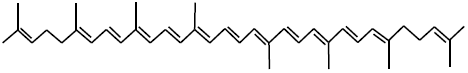
Structure of the C_40_ carotenoid Lycopene [97].

More than 750 carotenoids have been characterized in nature thus far, and these secondary metabolites are widely distributed among several biological taxa, including plants, fungi, and bacteria [56]. Carotenoids play essential roles in photosynthesis and photoprotection. In addition, these molecules are essential components of mammalian diets and have vital antioxidant activity [100, 101]. Although carotenoids are not typically characterized as having great pharmacological potential, they have attracted interest as important nutraceuticals due to their significant antioxidant and anti-cancer activities [57, 58]. Nutraceuticals are dietary or supplemental molecules that confer a physiological benefit, or provide biological protection against chronic conditions or diseases. Marine sponges are a source of such potential nutraceutical novel carotenoids.

Carotenoids have been also shown to provide both photoprotective and reproductive benefits in a variety of marine organisms [59, 60, 61]. Carotenoids play their vital photoprotective and antioxidant roles via high-light energy dissipation, and free radical detoxification. These secondary metabolites perform similar functions in marine sponges, and in particular shallow water or intertidal species where periodic exposure to excessive light and harmful UV irradiation occurs. Previous studies by Green and Koslovoa (1992) of sponges of the genus *Lubomirskia* in Lake Baikal show that carotenoid concentrations decrease as depth increases (from 2m to 17m). In addition, recent manipulative field studies of the common intertidal sponge *Clathria (Microciona) prolifera* from Chesapeake Bay, Virginia show carotenoid concentrations significantly decrease when the sponges were transplanted from light-exposed to shaded habitats; these data also point to the photoprotective function of carotenoids against harmful solar radiation [62]. It has been hypothesized that due to this photoprotective ability, carotenoids may play a central role in governing the ecological distribution of sponges such as *C. prolifera*, as these secondary metabolites would enable these organisms to live in what would otherwise be physiologically highly stressful environments [54, 61].

It is notable that, although unable to synthesize carotenoids *de novo*, marine sponges, particularly among the poecilosclerids and axinellids, are able to sequester these compounds in high concentrations [63, 64]. It has been postulated that in addition to diet, microbial and fungal symbionts are important sources of sponge natural products [65, 66]. Carotenoids found associated with sponges include aryl carotenoids, such as isorenieratene, renieratene, and renierapurpurin [67, 68], and except for sea sponges, these compounds had only been previously found in green sulfur bacteria [69, 67]. Thus it has been posited that symbiotic bacteria are the original source of these aryl compounds in sponges [67, 68]. In addition, acetylenic carotenoid sulfates termed bastaxanthins, and other related compounds thought to be fucoxanthin metabolites derived from microalgae, have been isolated from the marine sponges *Ianthella basta* [98] and *Prianos osiros* [70]. Recent molecular studies aimed at elucidating the main elements of the retinoid metabolic pathway in the demosponge *S. domuncula* revealed that the retinoid precursor β-carotene, which is enzymatically cleaved via the enzyme β-β-carotene 15,15’-dioxygenease to generate retinal, is generated by bacteria that form a symbiotic association with this sponge [71, 72].

### ii. Examples of Pharmaceutical Agents

#### Hemiasterlin

Hemiasterlins are found as secondary metabolites in several species of marine sponge (Cymbastela sp., Hemiasterella minor, Siphonochalina sp., and Auletta sp.) (9x). They are small cytotoxic tripeptides that disrupt the formation of microtubules in eukaryotic cells, via inhibition of tubulin polymerization. Both formation of the eukaryoyic cytoskeleton, and of the mitotic spindle, are disrupted, inducing mitotic arrest and inhibiting cellular proliferation. The mechanism by which hemiasterlin causes these effects is via drug binding to the vinca peptide site on the tubulin monomer (10x). Antimicrotubule targeting agents are among the most promising of anti-cancer agents due to their ability to effectively treat a wide variety of cancers (11x).

Existing antimicrotubule targeting agents are chemically diverse and generally act by either inhibiting or stabilizing the polymerization of tubulin, via binding to a one of a variety of binding sites on the tubulin surface. The vinca binding site sits at the polymerization interface of two tubulin monomers, and is targeted by the hemiasterlins. Other competitive binders to this site include dolastatins and the vinca alkaloids. Colchicine, taxanes and combretastatin are also antimicrotubule targeting agents, which act by binding to alternate, non-competitive sites on the tubulin subunit surface.

The hemiasteralin tripeptide consists of three highly substituted, unusual amino acids, and the structure is rigid. It is biosynthesized by non-ribosomal peptide synthase (NRPS).

The total synthesis of this molecule as well as several derivatives, has been done (12x, 13x, 14x). A naturally-derived analogue with higher activity has been designed, HTI-286. The terminal purine group of hemiasterlin is substituted by a phenyl group in this analogue, and the stereocenter of the adjacent bridging carbon is inverted, making the affinity of this molecule for its receptor higher than that of the lead compound.

Figure 4a. Cartoon of binding sites on the tubulin molecule. Kindly provided by Dr. Arie Zask, Columbia University and Wyeth Research, adapted from Dowling, Ann Rev Cell Dev Biol, 14, 89-111 (2000)

**Figure 4.**
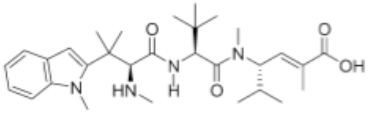
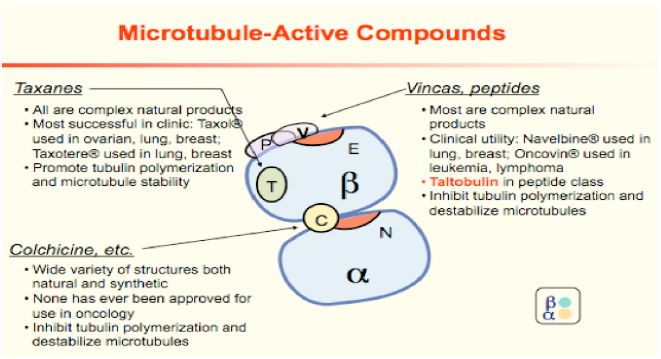
Hemiasterlin (from reference 11x)

#### Eribulin

Eribulin is an anticancer drug that treats certain forms of metastatic breast cancer, and may be used off-label for certain solid tumors, including those from prostate cancer, and non-small lung cancer tumors. It is a synthetic analogue [73] of the natural product halichondrin B, found in a variety of marine sponges, including *Axinella sp*., *Phakellia carteri Lissondendryx sp*. and *Halichondria okadai sp*. Originally discovered by Hirata and Uemura from the Meijo University in Nagoya, Japan in the waters of Miura Penisula, south of Tokyo, it was isolated using standard methods of natural product extraction. Briefly, the specimen is frozen, crushed and blended to a homogenate, followed by a series of by organic solvent extractions [74, 75].

Eribulin is a macrocyclic ketonic polyether macrolide [76]. Both eribulin and its parent compound halichondrin B are antimicrotubule agents whose mechanism of action is binding to a unique binding site on the plus end of the tubulin polymer (not monomer), inhibiting further microtubule growth via continued polymerization. Apoptosis is triggered after this blockade, resulting from the inhibition of proper tubulin formation within the dividing or growing cell.

**Figure 5.**
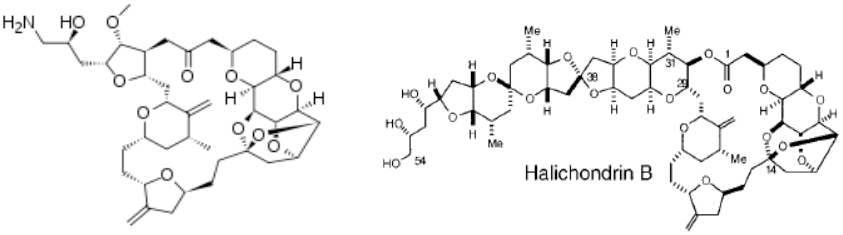
Eribulin and halichondron B (images courtesy of Wikipedia commons).

#### Ara-C (cytarabine)

Ara-C (cytarabine) is an aranucleoside, first discovered in 1945 by Warner Bergmann [76] in the Florida Keys, and isolated from the marine sponge Cryptotethia crypta sp. Originally termed a “spongonucleoside” this molecule acts as an effective and toxic anti-cancer drug, targeting cancer cells due to their rapid production of DNA as the cancer cells proliferate.

It has been noted that the only marine sponge found so far to produce free arabinosated nucleosides is Cryptotethia crypta. This sponge lives as a single organism (i.e., not as part of a coral reef or in any other external symbiotic relationship) and in shallow waters, partially submerged at times in sand. To prevent the entrance of sand into its pores as it filter-feeds, this species of sponge has the smallest known pores, which limits its intake of food both in flow rate and in total mass. The conjecture [77] is that this organism evolved a unique defense, arabinosated nucleosides, to defend against predators, as it is both vulnerable, living alone in the shallows and partly on land (sand), and only has access to a limited food supply which it must protect. This conjecture remains unproven but is compelling.

Aranucleosides are in the class of antineoplastic drugs known as antimetabolites. The sugar in these nucleoside inhibitors is arabinose instead of deoxyribose. Arabinose is structurally related to ribose, having a trans conformation of the hydroxyl groups on the C3 and C2 carbons of the sugar ring instead of syn. This class of drug acts by interfering with or blocking DNA replication and synthesis via blocking DNA polymerase [76].

Cytarabine remains the drug of choice for treatment of Hodgkin’s lymphoma, acute lymphoblastic leukemia and chronic myelogenous leukemia [78].

**Figure 6.**
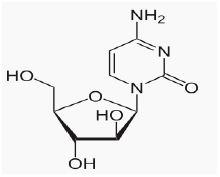
Cytarabine (image courtesy of Wikipedia commons)

#### Calicheamicin

Calicheamicins are produced by *Micromonospora echinospora spp calichensis* (NRRL l5839), a sapyphoritic actinomycete bacterium originally discovered in a chalky, limestone soil of Texas. Many actinomycetes are also found in marine environments and although this subspecies of *Micromonaspora* has not been explicitly found in the sponge biome, it may easily be a part of the symbiotic fauna found associated with that biome. Fermentation of organisms from the original soil sample, and fractionation of the products yielded showed anti-cancer activity due to calicheamicins on murine tumor models P388 and B16. Synthesis of calicheamicin was reported in 1996 (7), however fermentation remains the most economical way to produce this potent pharmacological molecule.

The calicheamicins are in the enediyene class of antitumor antibiotics, which contains some of the most potent antitumor agents ever discovered. The configuration of the pendant aryltetrasaccharide moiety allows specific targeting of the DNA minor groove (8) and subsequent strand scission of the DNA deoxyribose backbone, via carbon-centered diradical hydrogen abstraction mediated by the enediyne moiety.

The calicheamicin core structure is unique among all enediynes found to date. It is the only one to use an aryltetrasaccharide moiety instead of an intercalating moiety to effect binding to the target DNA. The enediyne group contains a bicyclic aglycone group along with an unusual and labile pendant trisulphide moiety that acts as a “trigger” which, upon its reduction, causes a modified Bergman cyclization resulting in the transformation of the enediyne group to a reactive diradical benzene intermediate which then attacks the DNA backbone, effecting a double-strand cleavage.

The structure and target specificity of this molecule implies that it may have evolved as a defense against foreign organisms, via cleavage of their DNA. Substitution of moieties of removal of any side group reduces or ablates the DNA-cleavage activity and specificity of this molecule.

**Figure 7.**
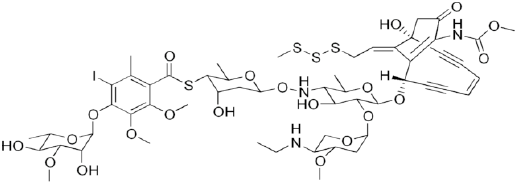
Calicheamicin (from Wikipedia Commons)

## 3. Conclusion and Future Aspects

The marine environment, a virtually untapped resource to date, holds great promise as a rich source of novel bioactive molecules, for discovery of both novel pharmaceutical agents and nutraceuticals, which have significant overlap with pharmaceuticals (as example, carotenoids can function as antioxidants, or as various potent cytotoxic agents).

Mining genomes to discover such novel, bioactive natural products is a powerful and effective discovery modality that is actively being developed by laboratories around the world. The goal is to conduct, as available, whole genome, proteome, transcriptome, lipidome, or metabolome analyses to discover novel marine bioactive, pharmaceutical, and nutraceutical agents, with the overall vision and goal of improving human health. Marine life and the marine environment, with its evolutionary diversity, wide range of ecological niches, dynamic self-organization of complex marine systems and microbial mats, and near-alien microenvironments (such as hydrothermal vents, and various and varied regions which include challenges to life such as hyperbaricity, hypersalinity, low photon penetration, and other challenges to evolution not found on the surface of this planet), is currently expected to be a rich source of such bioactive molecules, and further research and discovery is supported by these and other past findings.

## Acknowledgements

This manuscript benefitted greatly from lengthy discussions with Professor George Ellestad and Dr. Arie Zask, both of Columbia University. RRM thanks G.E. and A.Z. for critical reading of this manuscript and helpful suggestions. RRM thanks Gordon C. Sleeper for valuable critical commentary.

